# DNA binding domain undergoes dynamic and selective protein–protein interactions to facilitate CTCF insulation

**DOI:** 10.1101/2022.05.15.491938

**Authors:** Zhou Rong, Tian Kai, Huang Jie, Duan Wenjia, Fu Hongye, Feng Ying, Wang Hui, Jiang Yongpeng, Li Yuanjun, Wang Rui, Hu Jiazhi, Ma Hanhui, Qi Zhi, Ji Xiong

## Abstract

CTCF is required for three-dimensional chromatin organization and a predominant insulator protein. However, its roles in insulating enhancers have not been fully explained in 3D nuclear organization. Here, we found that the CTCF DNA binding domain (DBD) forms dynamic self-interacting clusters. We next investigated the spatial relationships between these clusters and other transcription regulators with a light-induced imaging system. Strikingly, CTCF DBD clusters were found to incorporate other insulator proteins but are not coenriched with transcriptional activators in the nucleus. This property is not observed in other domains of CTCF or the DBDs of other transcription factors. Moreover, endogenous CTCF shows a phenotype consistent with the DBD by forming small protein clusters and interacts less transcriptional activators bound, CTCF motif arrays. Our results reveal an interesting phenomenon that CTCF DBD interacts with insulator proteins and selectively localizes to nuclear positions with lower concentrations of transcriptional activators, providing new insights into the insulation function of CTCF.

## INTRODUCTION

Insulators are cis-regulatory elements that play a central role in regulating cell-type-specific gene expression during development and diseases(Flavahan et al., 2016; Herold et al., 2012). The insulation function blocks enhancer-activating promoters (Heger and Wiehe, 2014; Raab and Kamakaka, 2010; Recillas-Targa et al., 2002), and many protein factors, such as CTCF, BRD2, CHD8, and DDX5, have been reported to bind insulator elements and perform insulation functions (Bell et al., 1999; Hsu et al., 2017; Ishihara et al., 2006; Yao et al., 2010). Previous studies have investigated the DNA sequences that are required for CTCF-mediated insulation (Guo et al., 2015; He et al., 2021; Huang et al., 2021; Jia et al., 2020; Wang et al., 2021). CTCF has known to form loops with Cohesin by loop extrusion. Enhancers localize within the CTCF loops, and could not activate the genes outside of the CTCF loops (Dowen et al., 2014; Hnisz et al., 2016b; Ji et al., 2016; Sun et al., 2019). However, it is still difficult to understand how these CTCF loops could physically block enhancers to activate gene expression outside of the loops in the three-dimensional nuclear organization. CTCF binds to CCCTC DNA motifs in the genome and interacts with insulator proteins, and the roles of CTCF protein domains in insulation have not been fully explained in mammals (Bell et al., 1999; Nora et al., 2017; Zuin et al., 2014).

CTCF comprises an N-terminal domain (NTD), 11 zinc fingers, and C-terminal intrinsically disordered regions (IDRs) (Cuartero et al., 2019; Fudenberg et al., 2016; Ghirlando and Felsenfeld, 2016; Merkenschlager and Odom, 2013; Ong and Corces, 2014; Rolf Ohlsson, 2001; Vietri Rudan and Hadjur, 2015; Zlatanova and Caiafa, 2009). The NTD of CTCF interacts with Cohesin in chromatin to organize 3D chromatin structures through loop extrusion (Li et al., 2020; Nora et al., 2019; Pugacheva et al., 2020). The zinc finger 1 and 10, and C-terminal domains of CTCF show RNA binding activities (Hansen et al., 2019; Saldana-Meyer et al., 2014; Saldana-Meyer et al., 2019), and zinc fingers 3-7 of CTCF constitute the DNA binding domain (DBD), which directly interacts with DNA (Hashimoto et al., 2017; Yin et al., 2017). The RNA binding domain mediates CTCF self-interactions and is essential for CTCF-organized 3D chromatin structures. Although each domain of CTCF has been intensively studied in 3D chromatin organization, it is still difficult to understand how these CTCF loops block enhancer functions in the three-dimensional nucleus.

By taking advantage of the optoDroplet system for detecting weak, dynamic, and transient protein–protein interactions, we found that the CTCF DBD both interacts with itself and selectively interacts with other insulator proteins but is not coenriched with transcriptional activators in the nucleus, which is a new finding distinct from the generally assumed role of CTCF in DNA binding. Similar properties were not observed in other domains of CTCF or the DBDs of other transcription factors. Super-resolution imaging and bioinformatic and insulator reporter assays showed that endogenous CTCF forms small protein clusters and that its binding sites in the genome contain CTCF motif arrays that are associated with a low abundance of transcriptional activators and are positively correlated with insulator activity. Overall, we provide experimental evidence to help establish a new framework accounting for the insulation functions of CTCF.

## RESULTS

### Examination of dynamic CTCF DBD self-interactions with the optoDroplet system and *in vitro*-purified proteins

Transcription factors comprise low complexity domains (LCDs) and DBDs. Previous studies have shown that transcription factors can form local concentrated hubs through weak, transitory, dynamic LCD-LCD interactions (Chong et al., 2018). We sought to investigate whether the DBD also mediates dynamic protein–protein interactions, which may provide different functional aspects of transcription factors. To this end, we performed an optoDroplet assay with CTCF DBD (CTCF zinc figure 3-7, which is reported to directly interact with DNA (Hashimoto et al., 2017; Yin et al., 2017), details see method section). The optoDroplet system is a previously developed light-inducible reporter system used to determine which protein domains are able to self-interact to form protein clusters in mammalian cells (Shin et al., 2017) (Figure 1A). The optoDroplet experiments suggested that the CTCF DBD formed protein clusters, which were recognized as individual spherical, droplet-like objects, indicating the self-interaction of the CTCF DBD in cells (Figure 1B). We also found that the optoDroplet CTCF NTD did not form clusters and that the C-terminal IDR and RNA binding domain (RBD) could form self-interacting protein clusters (Saldana-Meyer et al., 2014) (Figure S1A), consistent with previous findings demonstrating that the disruption of the RBD affects CTCF clustering (Hansen et al., 2019; Saldana-Meyer et al., 2019). The IDR and RBD domains show the potential to form protein clusters (Hansen et al., 2020; Hansen et al., 2019; Saldana-Meyer et al., 2014; Saldana-Meyer et al., 2019), but the formation of protein clusters by the CTCF DBD was unexpected. Thus, we believe that the CTCF DBD that forms protein clusters is novel, and we focused on the CTCF DBD in the rest of this study.

**Figure 1.**
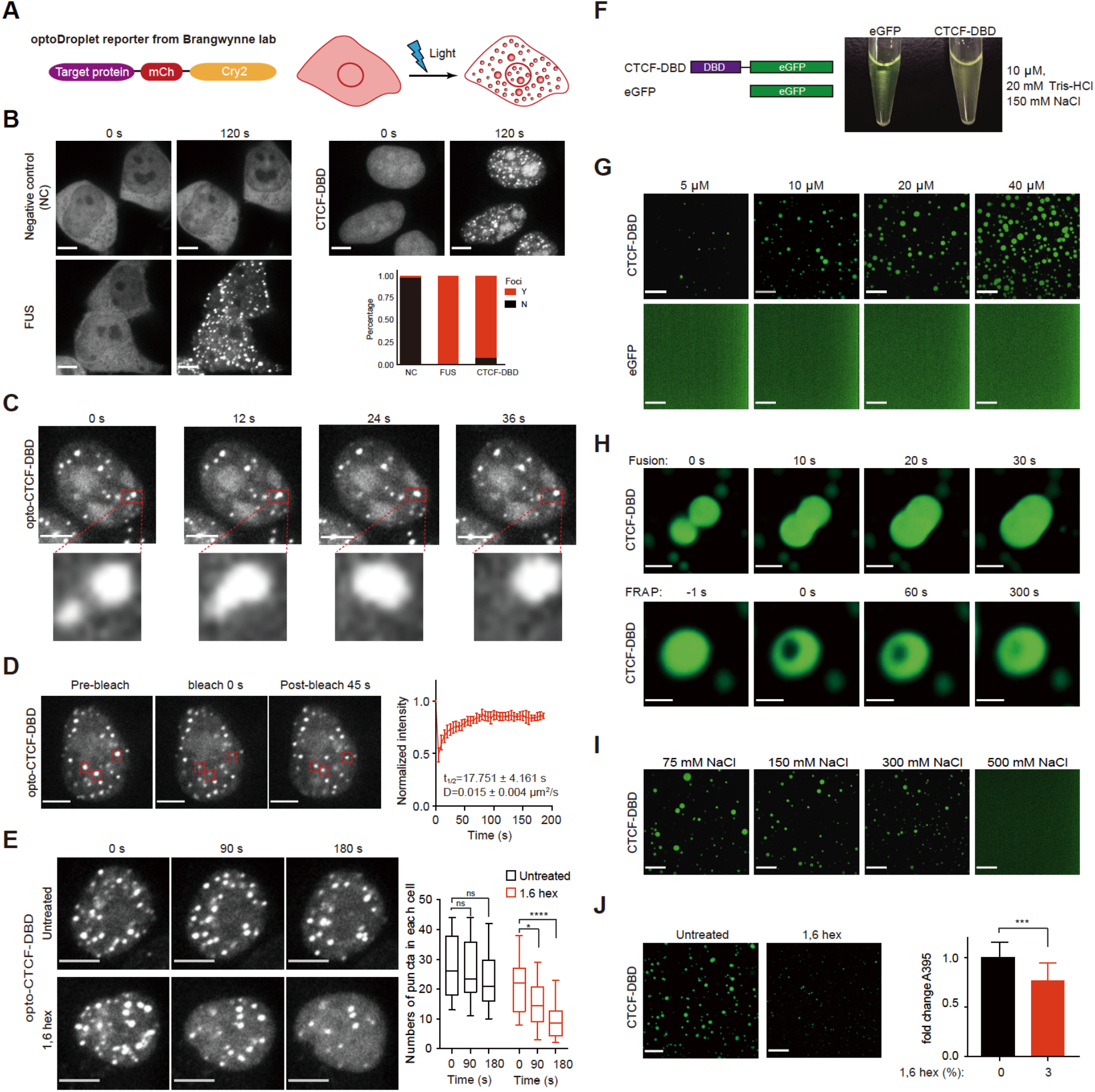
CTCF DBD undergoes self-interaction *in vitro* and in cells with optoDroplet. A. Schematic illustration of the optoDroplet reporter (left) and blue light-induced target protein domain clustering in live cells (right). B. Images of HEK293T cells expressing mCherry-Cry2, FUSN or the DBD of CTCF fused to mCherry-Cry2 (opto). Representative images of light-activated cells are shown. Fluorescent proteins expressed at similar levels were activated under identical conditions. The percentages of cells forming protein clusters are shown in the bar graphs. Y indicates observed clusters; N indicates that no clusters were observed. At least n=52 cells were used for the calculation. Scale bars, 5 μm. C. A CTCF DBD-opto cluster fusion event is shown, with a higher-resolution image below. Scale bars, 5 μm. D. Representative images of a FRAP experiment with CTCF DBD-opto in HEK293T cells. Red boxes indicate bleached clusters (left). Quantitative analyses of FRAP data of CTCF DBD-opto cluster (right). A bleaching event occurred at t=0 s. Data were plotted as the mean +/- SD (n=5). Scale bars, 5 μm, Apparent D: apparent diffusion coefficient; t1/2: half-time of recovery. E. Representative images of CTCF DBD-opto in HEK293T cells treated with 3% 1,6-hexanediol for 90 s and 180 s (left). Box plot illustration of the fold change in the number of CTCF DBD-opto clusters under 1,6-hexanediol treatment (right). n=20 in the control and n=20 in the 1,6-hexanediol treatment group were used for calculation. P values were calculated using the unpaired two-tailed Student’s t test (ns not significant, *<0.05, ****<0.0001). Scale bars, 5 μm. F. Schematic illustration of the recombinant EGFP and CTCF DBD-EGFP proteins used here (left). Turbidity analyses of the CTCF DBD and EGFP in buffer (20 mM Tris-HCl, 150 mM NaCl) at a concentration of 10 µM at room temperature (right). G. Representative images of droplet formation in the presence of different protein concentrations: CTCF DBD-EGFP or EGFP (bottom). Scale bars, 24 μm. H. Representative images of droplet fusion events and photobleaching recovery at the indicated time points. Scale bars, 5 μm (Fusion)/2.5 μm (FRAP). I. Representative images of CTCF DBD-EGFP droplet formation in the presence of different concentrations of NaCl. Scale bars, 24 μm. J. Representative images of the CTCF DBD droplet formation after treatment with 1,6-hexanediol (left) and absorbance analyses at 395 nm (A395) of CTCF DBD proteins in phase separation buffer (bottom right). P values were calculated using an unpaired two-tailed Student’s t test (***<0.001, **<0.01, *<0.05). Scale bars, 24 μm.

We next determined whether the optoDroplet CTCF DBD *per se* could form biomolecular condensates in cells. The features of condensates typically include a capacity for fusion, dynamic exchange with the local environment, and sensitivity to the disruption of hydrophobic interactions (Alberti et al., 2019; Shin and Brangwynne, 2017). Taking advantage of the optoDroplet system again, we monitored the fusion of CTCF DBD clusters in detail (Figure 1C, Supplementary Video 1). Our fluorescence recovery after photobleaching (FRAP) experiments indicated that the signals of the CTCF DBD clusters recovered within seconds upon photobleaching, similar to the recovery of previously reported condensates (Figure 1D, Supplementary Video 2). The FRAP recovery of the optoDroplet CTCF DBD was incomplete at 60 seconds and not fully recover after 200 seconds, suggesting that there might be different fractions of optoDroplet CTCF DBD in cells, as CTCF shows both bound and unbound fractions, and the dynamics of these two fractions are quite different (Soochit et al., 2021). Moreover, the CTCF DBD clusters were sensitive to 1,6-hexanediol; as treatment with this compound caused their dissolution within approximately 3 min (Figure 1E, Supplementary Video 3a-b). Collectively, these results indicate that the optoDroplet CTCF DBD exhibits condensate-like characteristics.

We examined whether the *in vitro*-purified CTCF DBD proteins exhibited biomolecular condensate features. We expressed and purified recombinant control EGFP and CTCF-DBD-EGFP fusion proteins to facilitate the detailed characterization of the cluster-forming behavior of the CTCF DBD (Figures S1B-D). The CTCF-RBD formed protein clusters, and CTCF-NTD did not form clusters using the similar experimental system (Figure S1E). Notably, the purified CTCF DBD fusion protein became opaque in buffer (20 mM Tris-HCl, 150 mM NaCl) at a concentration of 10 µM at room temperature, but purified EGFP did not (Figure 1F). Fluorescence microscopy analyses indicated that the CTCF DBD protein formed spherical clusters in a concentration-dependent manner, while EGFP did not (Figure 1G, Supplementary Video 4). Moreover, the CTCF DBD clusters could fuse and recover rapidly after photobleaching (Figure 1H, Supplementary Video 5) and were highly sensitive to the high-salt and 1,6-hexanediol treatments used to assess the phase separation behavior of the proteins (Figures 1I-J). These results collectively suggested that the *in vitro* protein–protein interactions of the CTCF DBD are weak, dynamic and transient, which is reminiscent of previously described LCD-LCD interactions (Chong et al., 2018).

### CTCF DBD optoDroplets selectively interact with insulator proteins and tend to avoid nuclear positions with a high abundance of transcriptional activators

Since the protein domains of CTCF can be efficiently clustered in cells with the optoDroplet system, we next sought to investigate how the domains of CTCF contribute to its protein interactions. We first cotransfected the CTCF DBD, RBD, NTD, or IDR optoDroplet plasmid with several eGFP-tagged transcriptional regulator plasmids. Then, the cells were exposed to blue light for up to 3 min to induce them to form relatively stable protein clusters, and the relative positions of the opto-fusions and appropriately expressed eGFP-fusions were analyzed. These experiments allowed us to examine many combinations of these sequences easily. The results showed that the insulator proteins BRD2 and CHD8 colocalized with CTCF DBD clusters but not with the CTCF RBD, NTD or IDR protein clusters after blue light induction (Figures 2A-C, S2A), which is consistent with previous observation that CTCF interacts with CHD8 and BRD2 (Bell et al., 1999; Hsu et al., 2017; Ishihara et al., 2006). Interestingly, BRD2 clusters appear to attract CTCF-DBD and CTCF DBD clusters seem to induce the clustering of CHD8, indicating that the behavior of BRD2 and CHD8 in relation to CTCF DBD is different (Figures 2A, 2D). The CTCF DBD clusters tended to be enriched at nuclear positions with lower concentrations of the transcriptional activators BRD3, OCT4, NANOG and SOX2, and these active apparatuses appeared to colocalize with RBD and IDR clusters (Figures 2A-C, S2A). GFP was distributed almost homogeneously in the nucleus with CTCF DBD clusters (Figure 2A) and served as a negative control.

**Figure 2.**
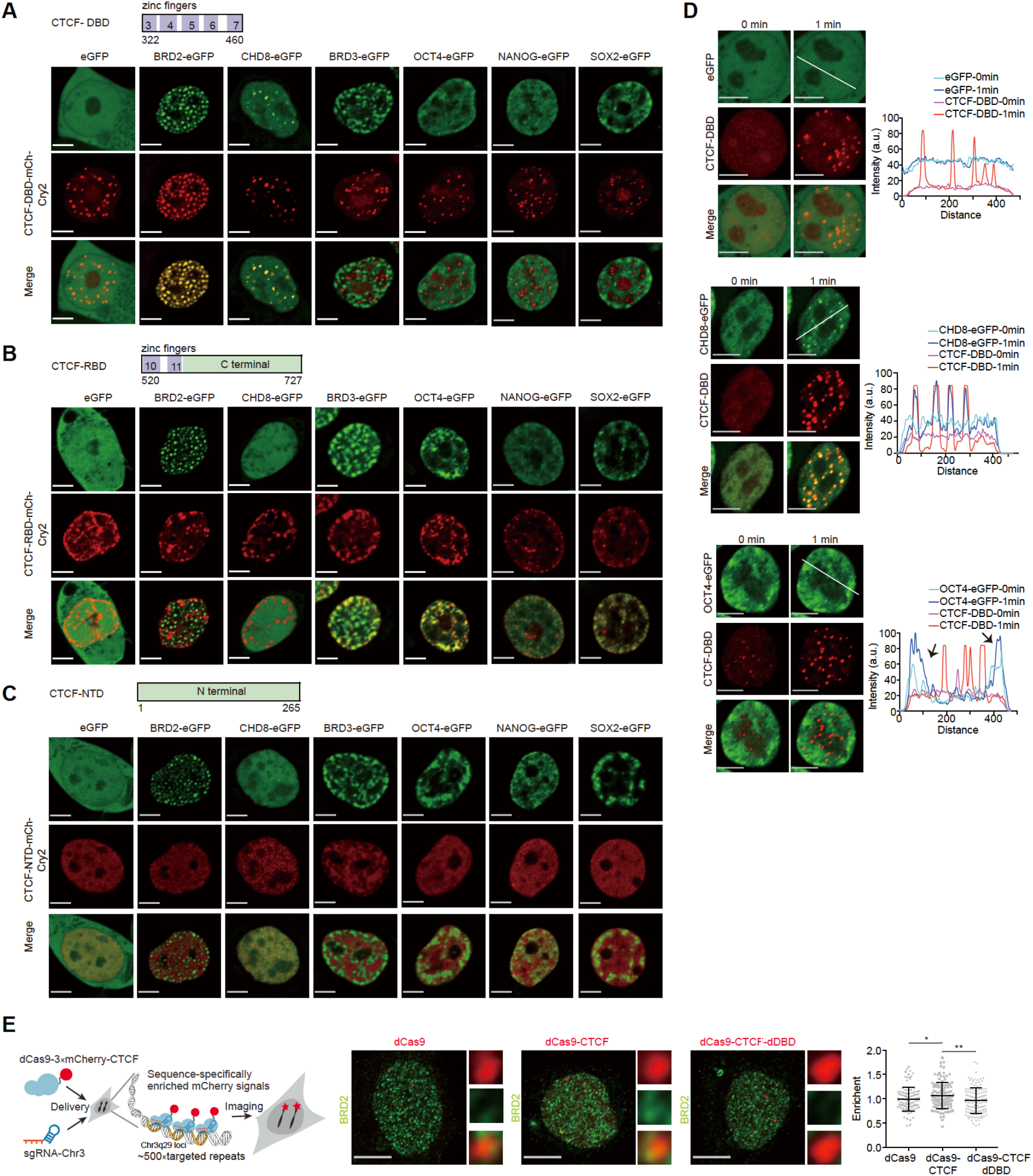
The CTCF DBD selectively interacts with insulator proteins but avoids transcriptional activators. A. Representative images of HEK293T cells expressing CTCF-DBD-mCh-Cry2 with eGFP, BRD2-eGFP, CHD8-eGFP, BRD3-eGFP, OCT4-eGFP, NANOG-eGFP, and SOX2-eGFP. Representative images of blue light-activated cells are shown. Scale bars, 5 μm. B. Representative images of HEK293T cells expressing CTCF-RBD-mCh-Cry2 with eGFP, BRD2-eGFP, CHD8-eGFP, BRD3-eGFP, OCT4-eGFP, NANOG-eGFP, and SOX2-eGFP. Representative images of blue light-activated cells are shown. Scale bars, 5 μm. C. Representative images of HEK293T cells expressing CTCF-NTD-mCh-Cry2 with eGFP, BRD2-eGFP, CHD8-eGFP, BRD3-eGFP, OCT4-eGFP, NANOG-eGFP, and SOX2-eGFP. Representative images of blue light-activated cells are shown. Scale bars, 5 μm. D. Fluorescence images of HEK293T cells expressing CTCF-DBD-mCh-Cry2 with eGFP (top), CHD8-eGFP (medium), and OCT4-eGFP (bottom) before and after 1 min of stimulation with blue light. Scale bars, 5 μm. The fluorescent intensity profiles at different positions in the CTCF DBD clusters before and after stimulation with light in the 488 nm and 561 nm channels (bottom) are indicated by a white line. Arrowheads indicate regions with increased signals after blue light stimulation. We observed that 0/19 CTCF DBD clusters excluded eGFP, 19/26 CTCF DBD clusters recruited CHD8-eGFP, and 5/20 CTCF DBD clusters excluded OCT4-eGFP. N/M = clusters showing exclusion or recruitment/total clusters. E. Left panel: the experimental design for dCas9 tethering of full-length CTCF to endogenous genomic loci. Middle panel: Representative immunofluorescence (IF) images of BRD2 in U2OS cells expressing sgRNA-ChrC3 with dCas9, dCas9-CTCF, or dCas9-CTCF-dDBD. The right images showed the magnified areas. Right panel: Scatter plot illustrating the enrichment of the intensity of the BRD2 immunofluorescence signal at the indicated tethered loci. At least n=66 cells were used for the calculation. Scale bars, 10 μm. P values were calculated using an unpaired two-tailed Student’s t test (**<0.01, *<0.05). Scale bars, 5 μm.

To provide evidence of the protein partner interactions of the CTCF DBD, we performed live-cell imaging before and after blue light irradiation for 1 min. The results showed that the CHD8 green fluorescent signals increased in regions within CTCF DBD optoDroplets, and the eGFP signals did not change (Figure 2D), indicating that our opto-quantification system functioned properly. The CTCF DBD optoDroplets appeared to converge at positions with a low density of OCT4 green fluorescent signals (Figure 2D). These results suggest that CTCF DBD protein clusters incorporate the insulator protein CHD8 and avoid positions with high densities of OCT4. Interestingly, not all CTCF DBD clusters occupied positions with a low density of OCT4 (Figure 2D), indicating that the avoidance behavior of the CTCF DBD is context dependent. Additionally, we used dCas9 to tether full-length CTCF and DBD-deleted CTCF to repeated genomic regions (Chen et al., 2013; Ma et al., 2015; Ma et al., 2016; Wang et al., 2018; Yao et al., 2010), such that the tethered proteins would be visible via fluorescent imaging. The results showed that tethering the CTCF increased BRD2 signals in the targeted regions, while tethering the CTCF with DBD deletion did not (Figure 2E). These results were consistent with the idea that CTCF undergoes protein-protein interactions via its DBD.

### Endogenous CTCF forms small protein clusters, interacts with CTCF motif arrays in the genome and avoids regions with a high density of transcriptional activators

We next performed CTCF immunofluorescence analysis using different antibodies and fixation protocols to enhance the protein signals. The results revealed that CTCF could indeed form nuclear clusters in mammalian cells (Figures S3A-B). The results of 3D structured illumination microscopy (SIM) imaging showed that endogenous CTCF formed small nuclear clusters in live cells (Figures 3A, S3C). The small clusters of CTCF overlapped with Cohesin, which was reported previously (Hansen et al., 2017). The endogenous CTCF protein clusters were also sensitive to 1,6-hexanediol treatment (Figure S3D). Real-time imaging of halo-tagged CTCF revealed a few small CTCF clusters in the early stage of mitotic exit, and these clusters subsequently grew larger as the cell cycle progressed (Figure S3E, Supplementary Video 6). These results are consistent with previously reported functions of CTCF and chromatin reorganization during mitotic exit (Abramo et al., 2019; Oomen et al., 2019; Zhang et al., 2019). The different reported distribution patterns of CTCF are due to the various imaging methods applied. As the endogenous CTCF clusters are very small, it is challenging to determine whether other protein factors colocalize with CTCF clusters or not through imaging technique. This is the reason that we used the optoDroplet system to enlarge the cluster signals, which are easily observed under low-resolution microscopy.

**Figure 3.**
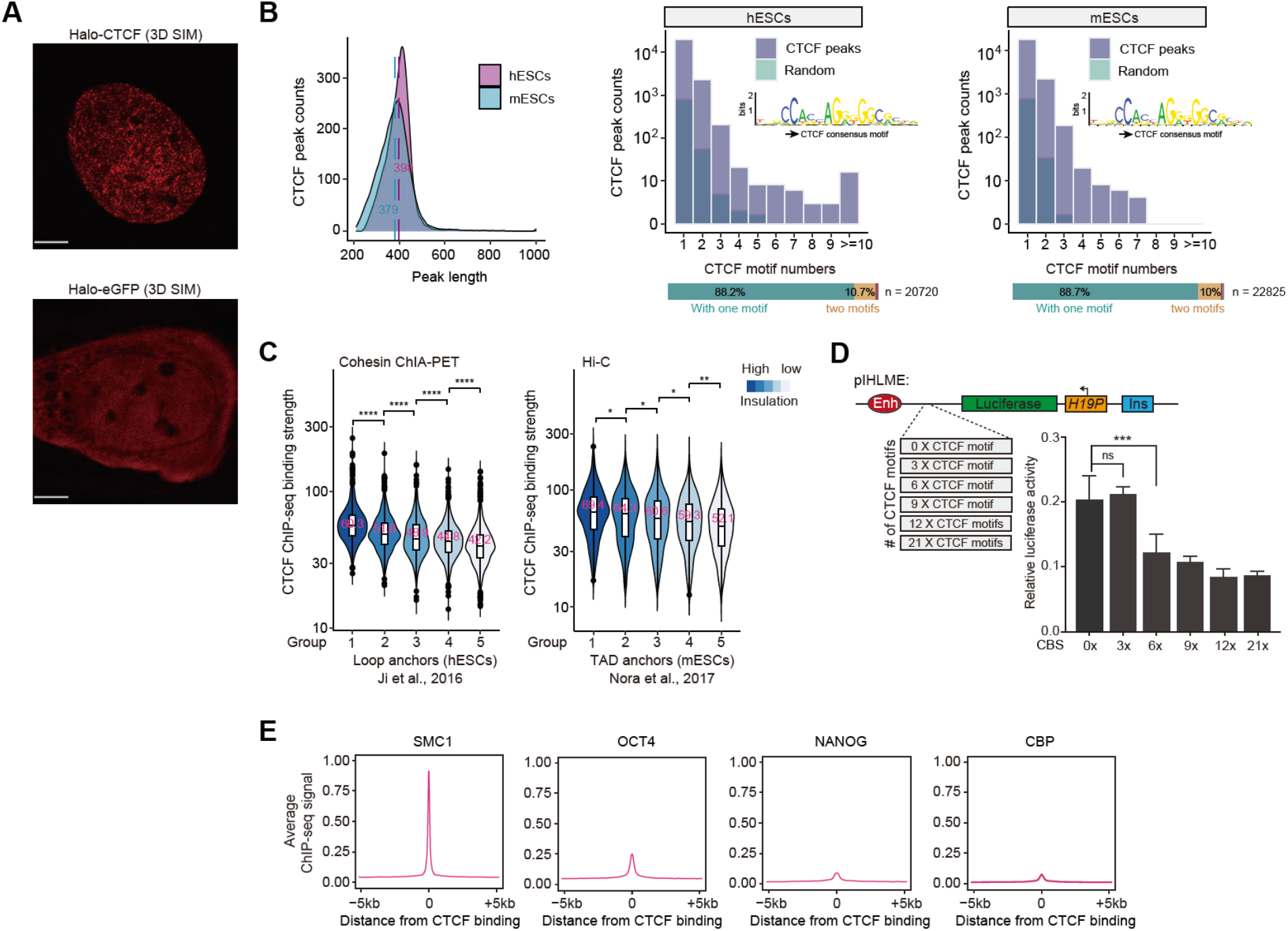
Full-length CTCF forms small protein clusters and interacts with CTCF motif arrays occupied by low densities of transcriptional activators. A. 3D SIM imaging of halo-eGFP (left) or halo-tagged CTCF (right) in U2OS cells. The red fluorescence signals indicate endogenous CTCF protein. Scale bars, 5 μm. Immunoblotting and genotyping identification results are shown on the right. B. Left: CTCF peak length distributions of ChIP-seq data generated from human embryonic stem cells (hESCs) and ChIP-exo data derived from mouse embryonic stem cells (mESCs). Right: Histogram displaying the CTCF motif occurrence in CTCF ChIP-seq peaks identified in hESCs and mESCs, with the bottom bar indicating the percentage of occurrence frequency of this motif. The light purple and cyan color refer to CTCF peaks and randomly selected regions in the corresponding human and mouse genomes. The position weight matrix for the canonical JASPAR CTCF motif finding is shown middle right. C. Left: Violin plot of CTCF ChIP-Seq signals corresponding to different insulation strengths at loop anchors from cohesin ChIA-PET data in hESCs (1-5: insulation strength from high to low). The analyses of the left anchor are shown, and the right anchor behaves in the same way. Right: Same as left but for TAD anchors from Hi-C data in mESCs. D. Schematic illustrations of the pIHLME reporter used in the luciferase assay (top). Luciferase activities of the pIHLME construct in the presence of the indicated number of CTCF motif insertions in HEK293T cells (bottom right). P values were calculated using two-tailed Student’s t tests (***<0.001, **<0.01, *<0.05). The CTCF motif sequences are shown in Supplementary Table 5. E. Averaged ChIP-Seq signals of cohesin (SMC1), OCT4, NANOG, and CBP at CTCF binding sites.

We sought to obtain functional insights about full-length CTCF in the genome through bioinformatic analyses. The number of CTCF motifs was calculated according to CTCF ChIP-Seq peaks derived from human embryonic stem cells (Ji et al., 2016), as these cells are normal cells cultured *in vitro,* and studies of these cells would reveal the normal functions of CTCF. More than 10% of CTCF-binding sites presented multiple CTCF motifs with regular ChIP and ChIP-exo datasets (Figure 3B, Supplementary Table 1), which is in accord with the idea that CTCF could execute its functions in chromatin through interactions with motif arrays (Schuijers et al., 2018). CTCF-CTCF loops have been shown to function as an insulated neighborhood (Dowen et al., 2014; Hnisz et al., 2016a; Ji et al., 2016). CTCF insulation scores were calculated by dividing the values of the Cohesin ChIA-PET signals within the loops by the values of the signals of the loops across loop anchors. The loop anchors were then subgrouped into 5 groups and ranked from high to low based on the insulation scores. The CTCF ChIP-Seq signals of each group were plotted in a violin diagram. The plot indicated that increased insulation scores for loop anchors were associated with stronger CTCF chromatin binding (Figure 3C). The same trend was also observed with Hi-C data (Figure 3C).

The pIHLIE luciferase reporter is widely used to evaluate the insulation activities of CTCF (Ishihara et al., 2006; Yao et al., 2010). The reporter consists of H19 promoter-driven firefly luciferase and an enhancer with a CTCF insulator between them. pIHLME is a version of pIHLIE with a mutated CTCF insulator sequence. Various numbers of CTCF motifs were inserted into the mutated regions of pIHLME, and a dual luciferase assay was performed. The analyses indicated that increasing the number of CTCF motifs resulted in a significant induction of insulator activity (Figure 3D). The CTCF binding site orientations were the same for all the CTCF motifs under our design, which may be the reason why the 3X CTCF motifs did not achieve insulation. We further investigated the interactions between CTCF and transcriptional activators in chromatin by performing colocalization analyses with published ChIP-Seq data. The signals of transcriptional activators (OCT4, NANOG and CBP) and Cohesin ChIP-Seq were plotted at the CTCF binding peaks. The analyses indicated that the CTCF binding peaks were associated with a high density of Cohesin binding, as expected, but showed little binding of OCT4, NANOG and CBP (Figure 3E). These results suggest that CTCF occupies regions with low densities of transcriptional activators in the genome. Note: this is just a correlative result and could not interoperate the causal effects.

### Arginine residues in the DBD are frequently mutated in various cancers and are critical for CTCF insulation

We next investigated the potential mechanism(s) of CTCF cluster formation via a variety of *in silico* analyses and mutational experiments. First, we noted that the CTCF zinc fingers were preferentially enriched with cysteine, histidine, and arginine residues (Figure 4A). The enrichment of cysteine and histidine residues was expected, as CTCF is a C2H2-type zinc finger transcription factor. The optoDroplet assay revealed that the samples bearing the variants with arginine mutations produced significantly fewer protein clusters than those with the wild-type CTCF DBD, so these mutations apparently generate CTCF variants with a reduced ability to form clusters, while cysteine and histidine appear to be necessary for the nuclear localization of the CTCF DBD (Figure S4A). Interestingly, an analysis of the COSMIC database using the entire CTCF open reading frame as a query indicated that the CTCF DBD is a cancer mutation hotspot (Figure 4B, Supplementary Table 2). Enrichment analyses of COSMIC mutations of the CTCF ORF across 24 different tissue types revealed that approximately 20% of CTCF mutations were associated with endometrioid carcinoma.

**Figure 4.**
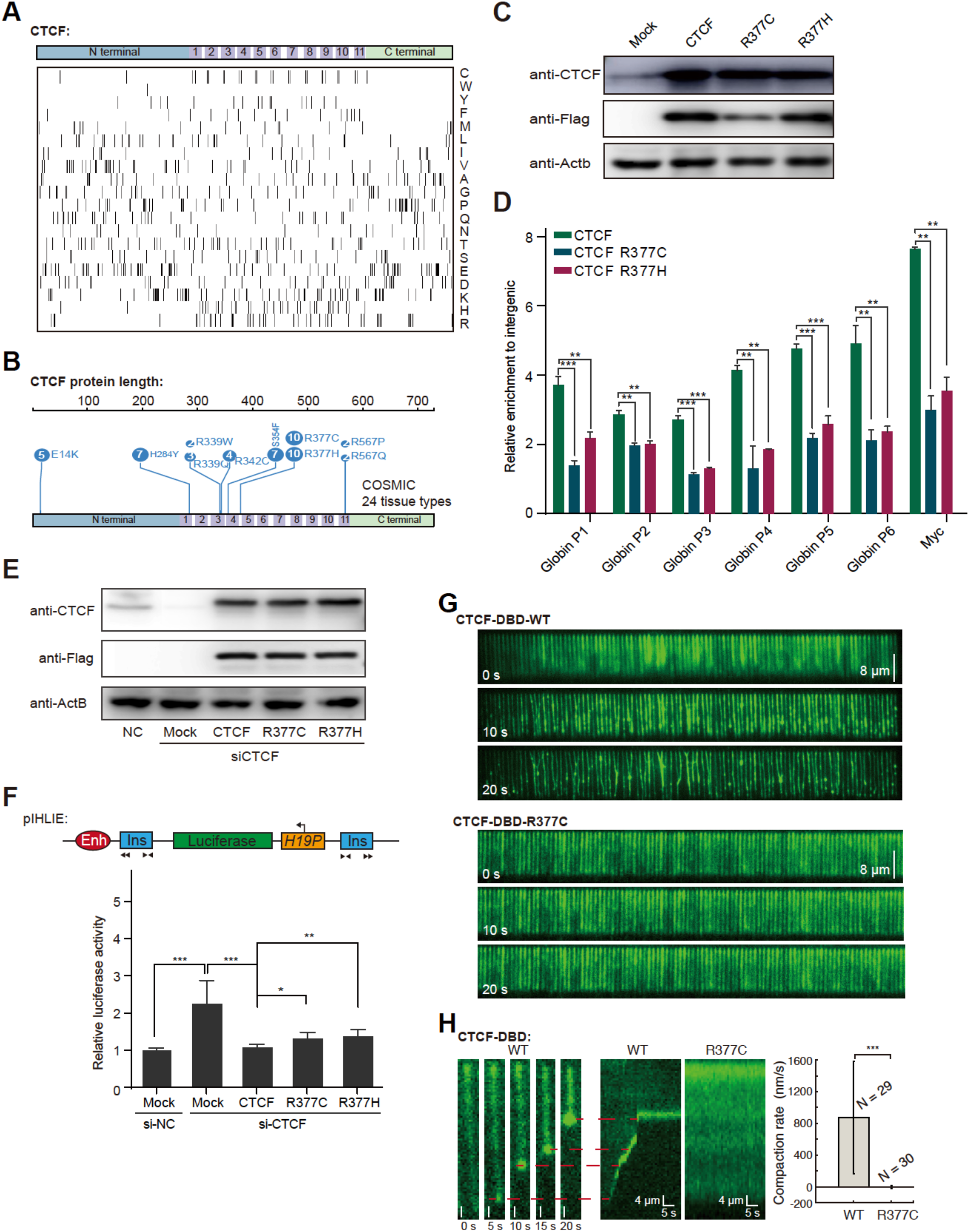
Arginine residues in the CTCF DBD are critical for self-interaction and insulation. A. Amino acid composition of the full-length CTCF protein. Black bars in each row indicate single amino acids, and the single-letter amino acid code is shown in the right panel. B. Somatic missense mutation hotspot landscape of the CTCF protein. Each number in the circle corresponds to the number of mutations at that amino acid position. The top 7 high-frequency mutation positions were chosen from the COSMIC database. C. Western blot of total cell lysates of HEK293T cells transfected with EGFP-fused full-length wild-type or cancer-associated mutant CTCF or the EGFP vector (mock). D. EGFP-fused full-length wild-type or cancer-associated mutant CTCF was transfected into HEK293T cells, and chromatin binding at C-MYC and b-globin loci was measured by eGFP ChIP–qPCR in transfected cells. Data are presented as the average of three replicates. P values were calculated using two-tailed Student’s t tests (***<0.001, **<0.01). The qPCR primer sequences are shown in Supplementary Table 5. E. Western blot of total cell lysates of HEK293T cells after the siRNA-mediated knockdown of CTCF and rescue with 2xFlag-fused full-length wild-type or cancer-associated mutant CTCF or the EGFP vector (mock). F. Schematic illustration of the pIHLIE reporter used in the luciferase assay (top). Luciferase activities of the pIHLIE construct in the presence of the empty vector (Mock), full-length CTCF, and cancer-derived mutants (R377C and R377H) in HEK293T cells. The luciferase signal was normalized to that of the internal control. Data are presented as the average of three replicates. P values were calculated using two-tailed Student’s t tests (***<0.001, **<0.01, *<0.05). G. Wide-field TIRFM image showing that the wild-type (left top) and R377C (left bottom) CTCF DBDs interacted with DNA at the indicated time points. H. Wide-field TIRFM images showing the compaction of a lambda DNA molecule with 1 μM wild-type CTCF DBD at each specific time point (left). Kymograph showing the compaction of the lambda DNA molecule with the wild-type or R377C CTCF DBD (right). Compaction rate of the wild-type (N = 29) and R377C CTCF DBDs (N = 30) (right bottom). The error bars represent the standard deviation (s.d.). The distribution was statistically compared using two-tailed Student’s t test (*** p < 0.001).

We generated variants of the 2 most frequently occurring DBD mutations (R377C and R377H) across 24 different tissue types in the COSMIC database. ChIP–qPCR analyses of the chromatin binding levels of the aforementioned CTCF variants for cancer mutations indicated that they showed significantly reduced binding at both *C-MYC* and *b-globin* loci (Figures 4C-D). The insulation activities of the CTCF mutants were then investigated with previously documented insulator reporters. The RNAi knockdown of CTCF decreased the insulator activity of the pIHLIE reporter, and the insulator activity could be rescued by the overexpression of an siRNA-resistant version of full-length CTCF. The overexpression of cancer-associated CTCF mutants partially rescued the insulator activities (Figures 4E-F). Together, the results indicated that the cancer-associated arginine mutations in the CTCF DBD interfered with the DNA binding and insulation functions of CTCF.

Previous studies have used the high-throughput single-molecule DNA curtain method (Zhao et al., 2017) to monitor interactions between DNA and heterochromatin protein 1α (HP1α) (Larson et al., 2017) and Vernalization 1 (VRN1) (Zhou et al., 2019). The authors of these studies indicated that the DNA “shrinking” behavior that they observed occurred due to a speculated liquid–liquid phase separation mechanism in DNA, suggesting a biological function of gene repression (Larson et al., 2017). We conducted similar experiments in which CTCF DBD was expressed and purified *in vitro*, and its DNA binding activity was confirmed (Figure S4B). DNA curtain analysis showed that the wild-type protein, but not the R377C mutant variant (from endometrioid carcinoma), could readily bind the DNA curtain, shrink DNA, and form bright fluorescent clusters at the ends of the DNA sequence (Figure 4G, Supplementary Video 7a-b). Specifically, the compaction rate of the wild-type CTCF DBD was measured as 871 ± 706 nm/s (mean ± s.d., Figure 4H). In comparison, the R377C variant did not shrink DNA under the investigated conditions (Figures 4G-H, Supplementary Video 8) but displayed a lower affinity of DNA binding when examined in electrophoretic mobility shift assays (EMSAs) (Figure S4C). The differences in DNA binding between the EMSA and DNA curtain experiments were due to the differences in these technologies. Our results collectively confirm the self-interactions of the CTCF DBD and indicate that a potentially pathogenically relevant mutation might compromise the clustering or DNA binding capacity of CTCF.

### More transcription factor DBDs exhibit selective protein–protein interactions

In light of our demonstration that the CTCF DBD clusters formed by self-interaction are capable of mediating the selective interactions of CTCF, we next explored whether this finding may be more generally applicable to other transcription factors. Hence, the optoDroplet assay was performed with two other C2H2-type DBDs (from BCL6 and YY1) and one GATA-type DBD (from GATA3), and the results showed that the DBDs of BCL6, YY1, and GATA3 each formed self-interacting protein clusters in HEK293T cells (Figure S4D). These results suggest that the DBDs of transcription factors may function via extensive protein– protein interactions, in addition to their intrinsic DNA binding activity. Our findings indicated that the optoDroplet CTCF DBD is capable of interacting with insulator proteins and avoiding high concentrations of transcriptional activators. A similar optoDroplet assay was performed with the BCL6 DBD and the GATA3 DBD. The results showed that BCL6-DBD and GATA3-DBD optoDroplets co-localized with the CHD8 and OCT4, but not for BRD2 and BRD3 (Figures 5A-B). The Figure 3A showed that CTCF DBD optoDroplets colocalized with BRD2 and CHD8, but not coenriched with the OCT4 and BRD3. These results suggest that the distribution relationships of BCL6 and GATA3 DBD optoDroplets with insulator proteins and transcriptional activators were different from those of CTCF DBD, implicating that the CTCF DBD may have different properties that other DBDs.

**Figure 5.**
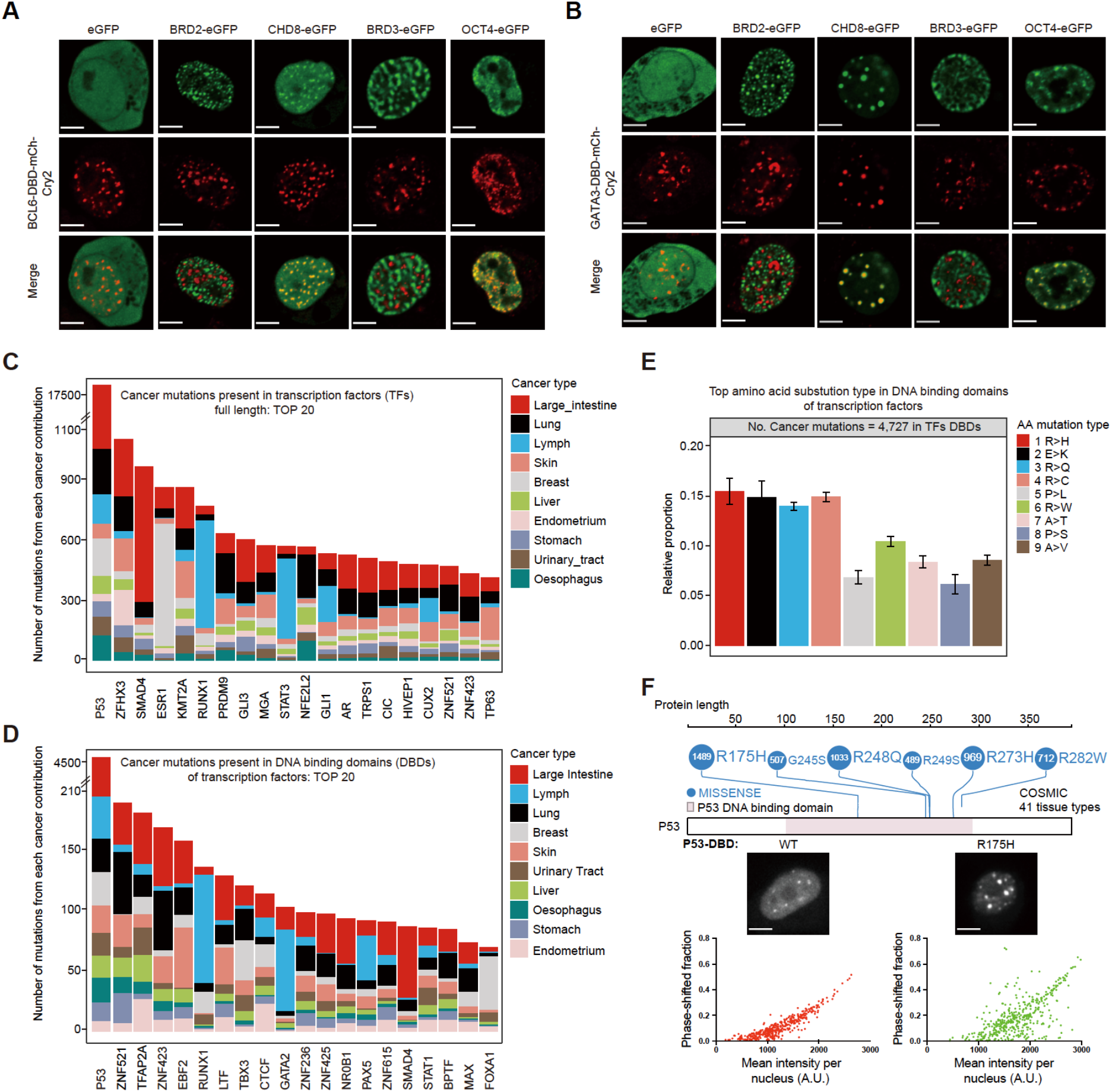
Additional transcription factor DBDs mediate selective protein– protein interactions. A. Representative images of HEK293T cells expressing BCL6-DBD-mCh-Cry2 with eGFP, BRD2-eGFP, CHD8-eGFP, BRD3-eGFP, and OCT4-eGFP. Representative images of blue light-activated cells are shown. Scale bars, 5 μm. B. Representative images of HEK293T cells expressing GATA3-DBD-mCh-Cry2 with eGFP, BRD2-eGFP, CHD8-eGFP, BRD3-eGFP, and OCT4-eGFP. Representative images of blue light-activated cells are shown. Scale bars, 5 μm. C. Top 20 transcription factors that are frequently mutated in cancers. The types of cancers are listed on the right. The most frequent types of missense amino acids in the DBDs of transcription factors in the COSMIC database are indicated as AA mutation types. D. Top 20 DBDs of transcription factors that are frequently mutated in cancers. The types of cancers are listed on the right. E. Most frequent types of missense amino acids in the DBDs of transcription factors in the COSMIC database. F. Hotspot somatic missense mutational landscape of the P53 protein. The top 6 high-frequency mutation positions were chosen from the COSMIC database (top). Images of the expression of the wild-type P53 (transcription factor) DBD (left bottom) and the P53 R157H cancer mutant DBD (right bottom) fused to mCherry-Cry2 in HEK293T cells. The percentage of the phase-shift fraction related to the mean fluorescence intensity per cell is shown with a scatter plot (bottom left), and the number of cells used for statistical analysis is shown in the plot. P values were calculated using the two-tailed Student’s t test (***<0.001, *<0.05). Scale bars, 5 μm.

Cancer-associated arginine mutations in the CTCF DBD interfered with its DNA binding and insulation functions, so we considered whether these findings might point toward a potentially widespread mechanism in cancer. By searching the unique gene IDs of transcription factors from the COSMIC database (v87), we identified 174,974 missense variants in 1,254 human transcription factors (Barrera et al., 2016; Deplancke et al., 2016; Lambert et al., 2018), and 501 transcription factors presented 23,313 total missense variants in their DBDs (Figures 5C-D, Supplementary Table 3-4). We noted that the large intestine, lymph nodes, and lungs were the 3 organs harboring the largest numbers of missense mutations in the DBDs of transcription factors (Figure 5D, Supplementary Table 4). The average relative contributions of the amino acid substitutions types of the mutations found in each cancer type were calculated, which showed that arginine was the most frequently mutated amino acid (R>H, R>Q, R>C and R>W, 4 of the 9 most frequent amino acid mutation types) among the COSMIC mutations in the full-length proteins and DBDs of transcription factors (Figures 5E, S4E). The single most frequently mutated transcription factor, P53, had 7,992 missense mutations in its DBD among 41 different cancers in the COSMIC database (Figure 5F, Supplementary Table 4). These results suggest that arginine residues are frequently mutated in cancers. We further cloned the wild-type DBD of P53 and a hotspot mutant (R175H) this DBD into the optoDroplet reporter. Surprisingly, the P53 DBD formed protein clusters, and the cancer-associated P53 DBD mutation obviously affected protein cluster formation in HEK293T cells (Figure 5F). Compared with CTCF DBD, the results imply that the arginine residues contribute differently to protein clustering for different transcription factors. This evidence is consistent with a potential mechanism in which cancer mutations in transcription factor DBDs result in the dysregulation of self-interactions.

## DISCUSSION

The CTCF/Cohesin complex-mediated loop extrusion model explained the molecular basis of 3D genome organization well, but it is still challenging to understand how CTCF/Cohesin chromatin loops block enhancer functions in three-dimensional nuclear organization. Specifically, we demonstrate that the DBD of CTCF undergoes dynamic self-interaction independent of its IDR *in vitro* and in cells with optoDroplet. The CTCF DBD selectively interacts with insulator proteins and avoids transcriptional activators. Other domains of CTCF do not exhibit similar properties. Accordingly, endogenous CTCF forms small protein clusters and binds genomic regions with a high abundance of CTCF motifs but low densities of transcriptional activators, consistent with a spatial segregation model of CTCF insulation (Figure S5A). Furthermore, the DBDs of other transcription factors show selective protein–protein interactions, and arginine residues are frequently mutated in various cancers. Our results reveal a previously underappreciated function of the DBD: the ability to engage in selective, dynamic and transient protein–protein interactions, which provides new insights for understanding transcription factor function in development and diseases. These CTCF DBD results inspired us to propose a spatial segregation model that CTCF selectively interacts with insulator proteins, and avoids nuclear positions with a high density of the transcriptional activators, which may spatially block the communication of the transcriptional apparatus of enhancers to active its targeted promoters (Figure S5A). This model was based on the results of CTCF DBD, which warrants further investigation of full-length endogenous CTCF in the future.

Recent studies have shown that CTCF forms small protein clusters in the nucleus, which is required for proper 3D chromatin organization and gene expression (Hansen et al., 2019; Saldana-Meyer et al., 2019). Recombinant purified CTCF and RNA form multimers of more than 2 megadaltons *in vitro* (Saldana-Meyer et al., 2014). Our current results could be simply explained by the liquid–liquid phase separation (LLPS) concept (Alberti et al., 2019; Brangwynne et al., 2009), but they are not sufficient to reach a conclusion about whether endogenous full-length CTCF forms LLPS or not, because our current evidence is mainly based on studying the CTCF DBD under artificial conditions. Recent quantification results showed that the endogenous concentration of CTCF is approximately 0.1 µM in mammalian cells (Cattoglio et al., 2019; Holzmann et al., 2019), while our *in vitro* droplet assay indicated that at a concentration of 10 µM, the CTCF DBD formed phase-separated condensates. We also found that the total CTCF protein levels could not be dramatically induced after transfection of exogenously expressed CTCF, suggesting a possible autoregulation mechanism of CTCF protein as previously observed (Kung et al., 2015). However, we cannot exclude the possibility that a high concentration of CTCF might result in phase separation of the protein under unusual circumstances, such as senescence, in specific cell types (Zirkel et al., 2018) or concentration of the protein at centrosomes during mitosis (Burke et al., 2005; Xiao et al., 2015). We also believe that our CTCF DBD-mediated protein clusters may play a role in the regulation of small clusters of endogenous CTCF at endogenous concentrations.

The cooperation among transcription factors is usually explained by transcription factor-transcription factor interactions, transcription factor-mediated DNA bending, or combinatorial interactions with the transcriptional machinery (Lambert et al., 2018; Levine et al., 2014; Martinez and Rao, 2012; Spitz and Furlong, 2012); however, the molecular basis of these interactions remains elusive. Here, we serendipitously found that the CTCF DBD undergoes self-interaction and mediates the formation protein clusters that incorporate insulator proteins but avoid transcriptional activators. The biophysical features of the CTCF DBD that we observed in this study are well correlated with the current knowledge of CTCF insulation, while other domains of CTCF do not show these features. The DBD clustering phenomenon is also likely to apply to many other transcription factors, which would provide new assays for visualizing the relationships between different transcription factors. The functions of full-length CTCF in cells are also variable, such CTCF occupies promoter-proximal regions for gene activation (Kubo et al., 2021; Schuijers et al., 2018), and CTCF binds to the distal insulator region, which blocks enhancers from activating gene promoters(Bartolomei, 2009; Hou et al., 2008). Besides this, CTCF has diverse functions from DNA replication, DNA repair, Pol II transcription, and splicing etc. (Chernukhin et al., 2007; Hwang et al., 2019; Lang et al., 2017; Shukla et al., 2011; Zhang et al., 2019; Zhao and Dean, 2004). We found different protein-protein interaction properties for the CTCF DBD and CTCF RBD, implicating that the biophysical properties of the different domains are variable. It is possible that different domains carry out different functions under different environmental conditions. This is also consistent with the diverse functions of CTCF in the nucleus.

We are just beginning to identify and investigate the activity of CTCF in avoiding transcriptional activators in cells, and standards for defining this avoidance activity have not yet been developed (McSwiggen et al., 2019). The observation that CTCF avoids regions with high densities of transcriptional activators raises more questions than it answers. For example, how does CTCF avoid transcriptional activators in the nucleus? There are approximately 1,200 transcription factors in humans (Lambert et al., 2018). How many of these transcription factors also form protein clusters, and do they also incorporate relevant factors and avoid opposing factors in three-dimensional nucleus? Importantly, the disruption of CTCF clustering has been shown to disrupt chromatin looping and dysregulate global gene expression (Hansen et al., 2019). We believe that DNA binding activity is essential for CTCF functions, but we would like to highlight that the protein–protein interactions mediated by the DBD may play an additional, important role. For the detailed molecular basis for the CTCF DBD-mediated protein-protein interactions and DNA are currently unknown. It could be: CTCF DBD interacts with other CTCF molecules while bound to DNA; or DNA bound CTCF interacts with other CTCF DBD through a different domain; or CTCF interacts with other CTCF when it is not bound to DNA. We believe the relationship with DNA binding is worthy to investigate in the future.

## Limitations of the study

This study revealed an unexpected clustering effect of CTCF DNA binding domains, and further showed the selective protein-protein interactions for CTCF DBD to facilitate CTCF-mediated insulation. However, whether and how the full-length CTCF exhibits similar property through it is DNA binding domain have not been documented in the current study. Besides, the functional relationship between CTCF DBD clustering and DNA binding capacity is also unclear. Therefore, further investigation on these two limitations would demystify the insulator functions of CTCF in mammalian cells.

## ACKNOWLEDGMENTS

We thank Drs. Clifford Brangwynne, Yanli Wang, Mitsuyoshi Nakao, Hongjie Yao, Yuanchao Xue, Jinyan Fu, and Richard Young for sharing plasmids and cell lines. We thank the members of the Ji laboratory for engaging in helpful discussions. This work was supported by funds from the Ministry of Science and Technology of China, the National Natural Science Foundation of China (Grants 2017YFA0506600, 31871309 and 32170569), and the Qidong-SLS Innovation Fund; this work was also supported by grants from the Peking-Tsinghua Center for Life Sciences and the Key Laboratory of Cell Proliferation and Differentiation of the Ministry of Education at the Peking University School of Life Sciences to X. J.; NSFC Grant No. 31670762 to Z.Q., and a grant from the China Postdoctoral Science Foundation to J. H (2017M610700).

## AUTHOR CONTRIBUTIONS

X.J. conceived and supervised the project. R.Z. performed the IF analyses, optoDroplet analyses, and *in vivo* and *in vitro* droplet analyses; W.J.D. performed the CTCF mutation analyses; K.T. performed all the luciferase analyses of CTCF insulator function. J.H. performed all the bioinformatic analyses. H.Y.F. and Q.Z. performed the EMSA and DNA curtain analyses. Y.F. and H.H.M. performed imaging analyses of endogenous CTCF during the cell cycle. T.K., H.W. and Y.P.J. performed the CTCF ChIP–qPCR analyses. Y.J.L. generated the subsets of constructs used for the optoDroplet experiments. All authors contributed to data analysis and data interpretation. X.J. wrote the manuscript with input from Z.Q., H.H.M., R.Z., K.T., and J.H. and help from the other authors.

---

**Figure S1.**
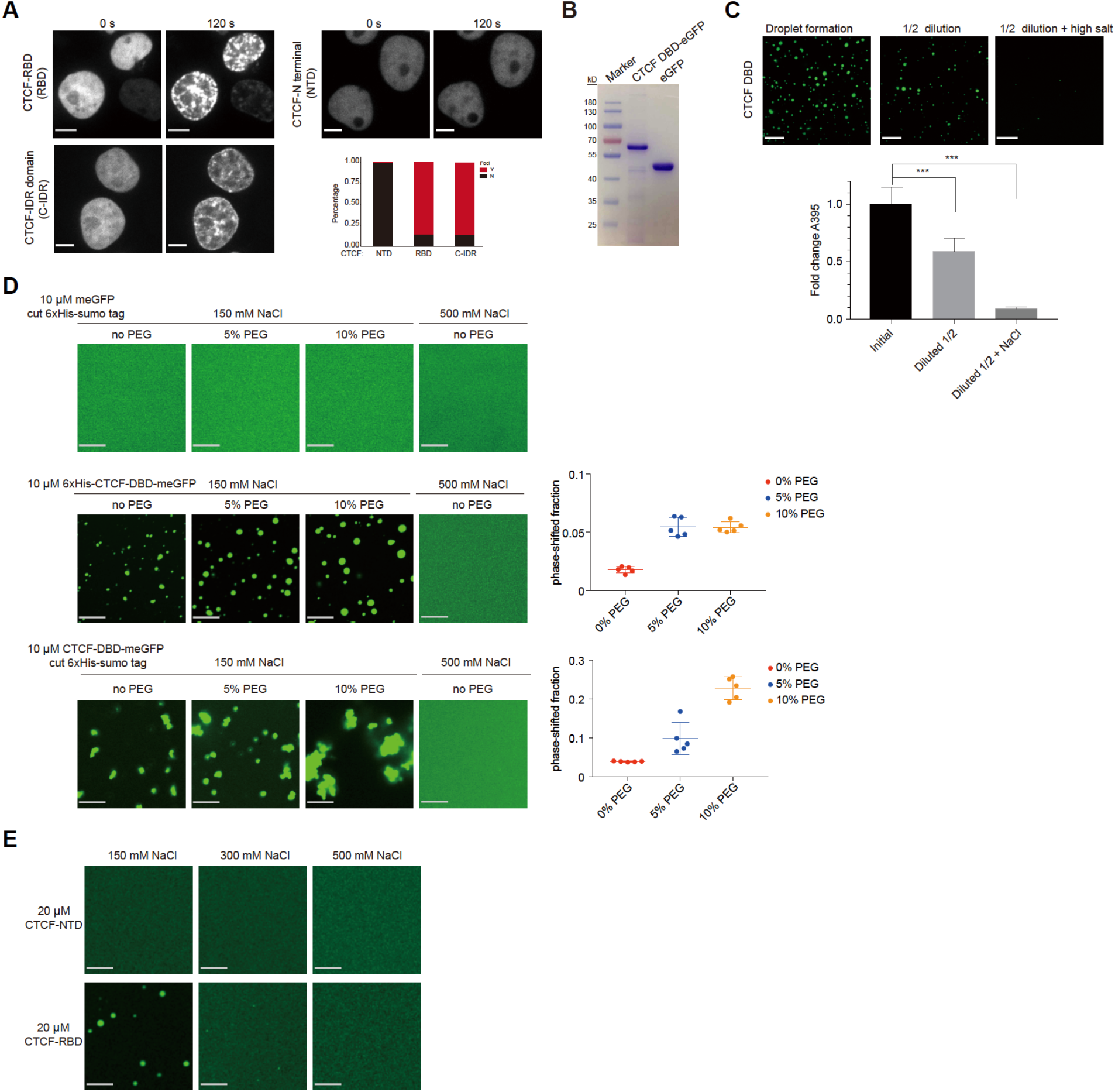
Self-interaction of the CTCF DBD *in vitro*. A. Images of HEK293T cells expressing the RNA binding domains of CTCF fused to mCherry-Cry2 (opto). Representative images of light-activated cells are shown. The fluorescent proteins expressed at similar levels were activated under identical conditions. The percentages of cells in which protein clusters are forming are shown in the bar graphs. Y indicates observed clusters; N indicates no observed clusters. At least n=52 cells were used for the calculation. Scale bars, 5 μm. B. SDS–PAGE followed by Coomassie blue staining analyses of purified recombinant CTCF DBD-EGFP and EGFPs. C. Representative images of the droplet reversibility assay. The CTCF DBD was tested with an initial buffer consisting of 10 mM protein and 150 mM NaCl at a 1:1 dilution (diluted ½) or high-salt buffer containing 500 mM NaCl. Scale bars, 24 μm. D. Representative images of 10 μM meGFP, 6xHis-DBD-meGFP (monomer form of eGFP), and DBD-meGFP droplet formation in the presence of different concentrations of PEG or NaCl. Scale bar, 10 μm. Quantification of the phase-shift fraction of 6xHis-DBD-meGFP and DBD-meGFP in the presence of different concentrations of PEG. E. Representative images of 20 μM CTCF NTD (top) and CTCF RBD (bottom) droplet formation in the presence of different concentrations of NaCl. Scale bars, 24 μm.

**Figure S2.**
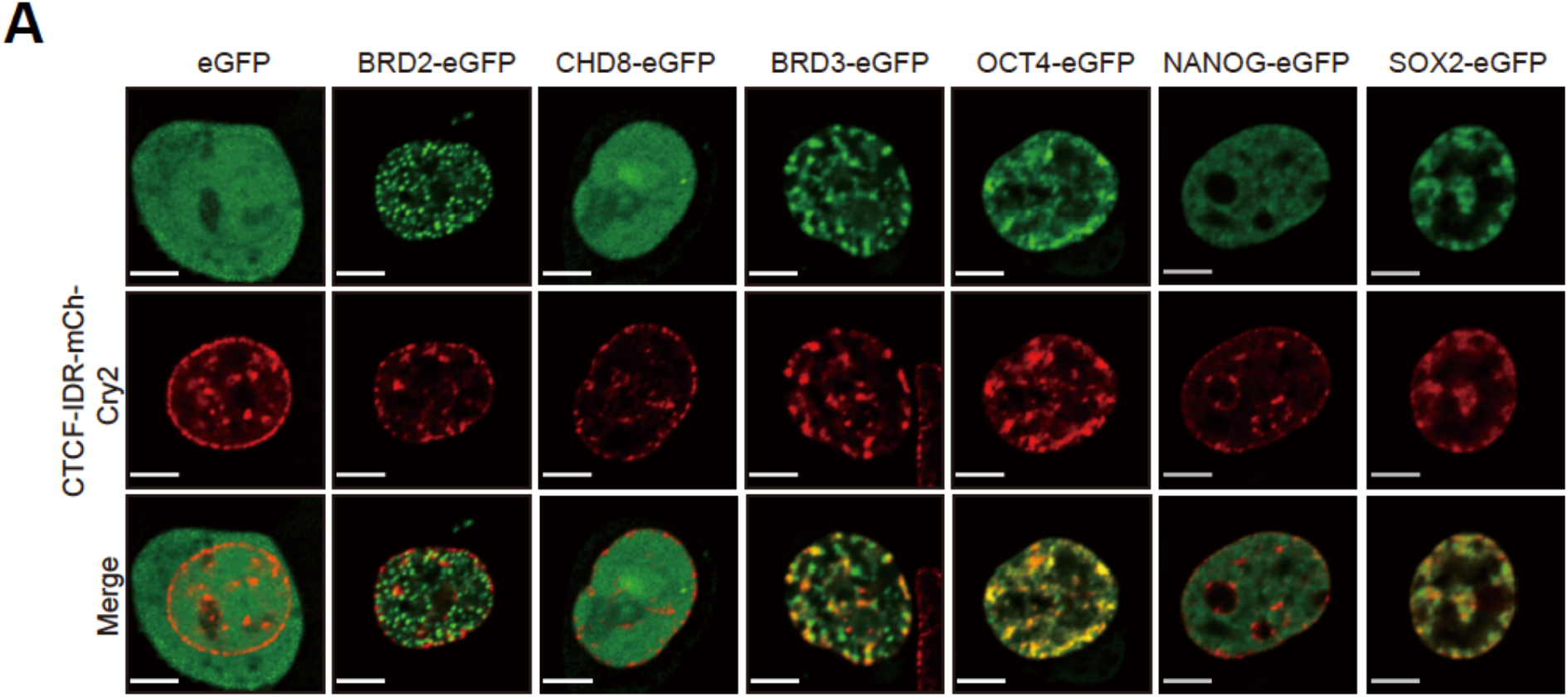
CTCF-IDR optoDrplets interact differently than the CTCF DBD. A. Representative images of HEK293T cells expressing CTCF-IDR-mCh-Cry2 with eGFP, BRD2-eGFP, CHD8-eGFP, BRD3-eGFP, OCT4-eGFP, NANOG-eGFP, and SOX2-eGFP. Representative images of blue light-activated cells are shown. Scale bars, 5 μm.

**Figure S3.**
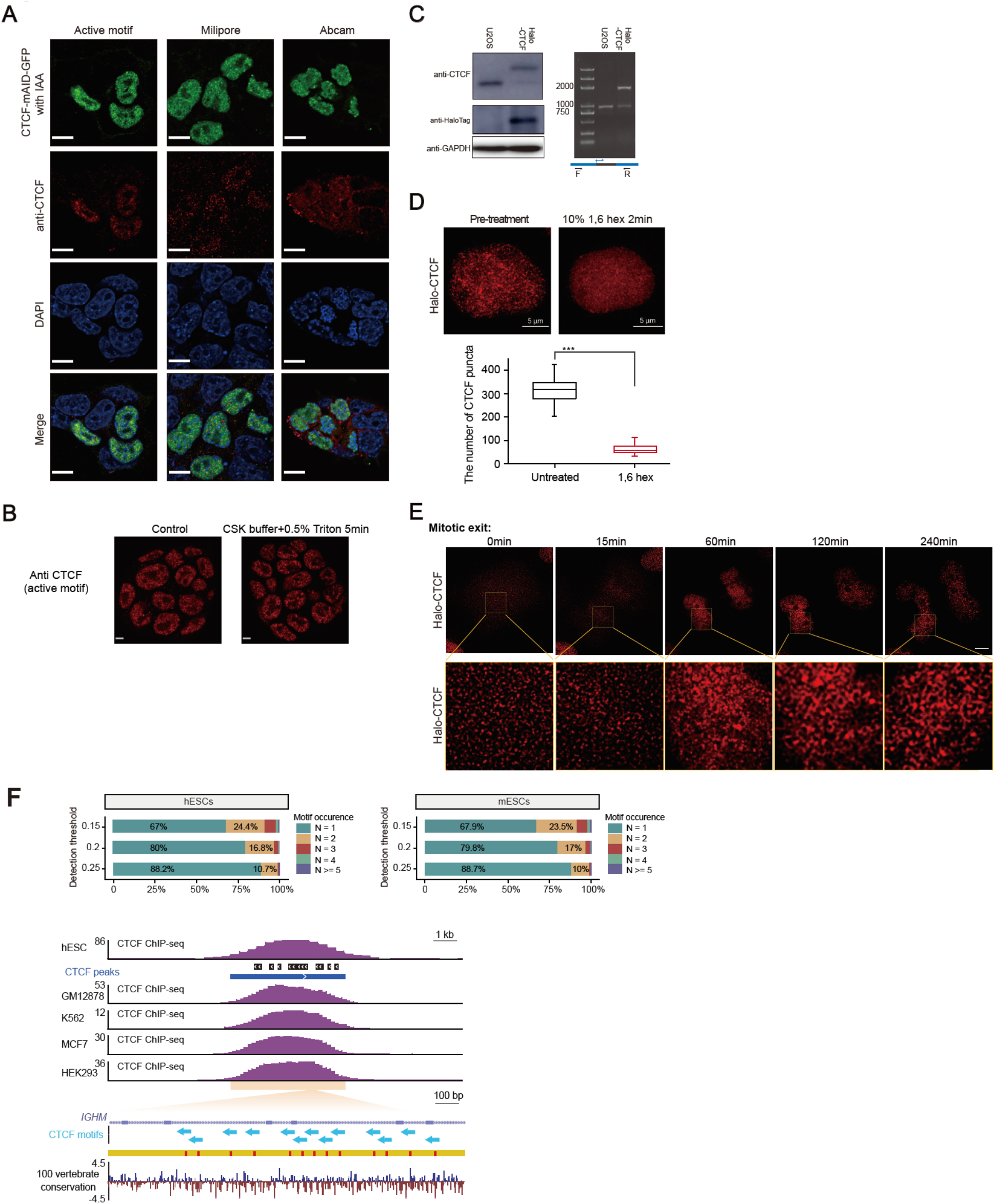
Endogenous CTCF forms small protein clusters. A. Immunofluorescence (IF) images of CTCF-mAID-GFP cells obtained using different antibodies. Endogenous CTCF-mAID-GFP is shown in green, the fluorescence signal is shown in red, and DAPI staining is shown in blue at the bottom. Scale bars, 10 μm. B. Immunofluorescence (IF) imaging of CTCF-mAID-GFP cells using different fixation protocols. Scale bars, 5 μm. C. Immunoblotting and genotyping identification of Halo-tagged CTCF in U2OS cells are shown. D. HaloTag-CTCF U2OS cells were imaged. HaloTag-CTCF was visualized by adding 200 nM HaloTag-JF549, and cells were chosen to track the formation of CTCF clusters during mitotic exit. The cells were imaged every 15 min for 240 min. Top: scale bars, 5 μm. Bottom: scale bars, 1 μm. E. Representative images of halo-tagged CTCF in U2OS cells under treatment with 10% 1,6-hexanediol for 2 min. Box plot illustrating the fold change in the number of CTCF halo-tagged CTCF clusters under 1,6-hexanediol treatment (bottom). n=68 in the control group and n=70 in the 1,6-hexanediol treatment group were used for the calculation. P values were calculated using an unpaired two-tailed Student’s t test (***<0.001, ****<0.0001). Scale bars, 10 μm. F. Upper: stacked bars representing the percentage of occurrence frequency of CTCF motif at different detection thresholds for CTCF peaks in hESCs and mESCs. Lower: Genome browser view of CTCF ChIP–seq signal across different cell lines around representative CTCF peak (IGHM, chr14:106,762,381-106,768,280, hg19). The middle panel depicts all putative CTCF binding motifs as blue arrows, which indicate the orientation of the motif. 100-way vertebrate conservation obtained from the UCSC genome browser is depicted at the bottom.

**Figure S4.**
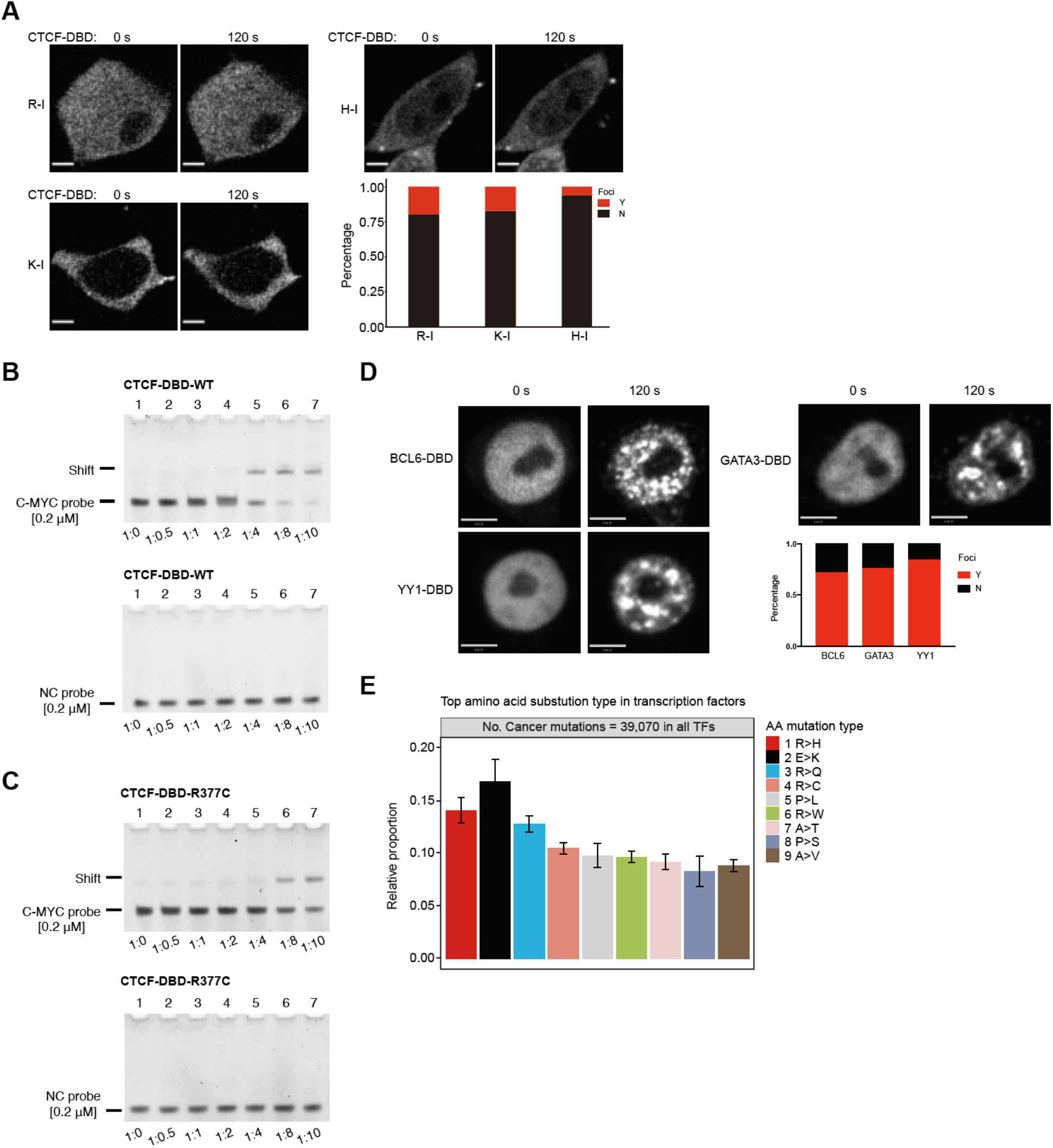
Arginine residues are critical for the DNA binding and self-interaction of the CTCF DBD. A. Representative images of light-activated HEK293T cells expressing CTCF DBD-saturated mutants (R-I, H-I, and K-I) fused to mCherry-Cry2. The plotted data came from at least n=37 cells under each condition. Scale bars, 2 μm. B. Electrophoretic mobility shift assay (EMSA). DNA (0.2 μM) (C-MYC target DNA (top) or a negative control (NC) DNA probe (bottom)) was incubated with the purified wild-type CTCF DBD and analyzed via native PAGE. C. Electrophoretic mobility shift assay (EMSA). DNA (0.2 μM) (C-MYC target DNA (top) or a negative control (NC) DNA probe (bottom)) was incubated with purified CTCF-DBD-R377C and analyzed via native PAGE. D. Representative images of HEK293T cells expressing the DBDs of BCL6-mCh-Cry2 (left) and GATA3-mCh-Cry2 (right) with BRD2-eGFP and OCT4-eGFP. Representative images of light-activated cells are shown. Scale bars, 5 μm. E. Most frequent types of missense amino acid mutations in the transcription factors in the COSMIC database.

**Figure S5.**
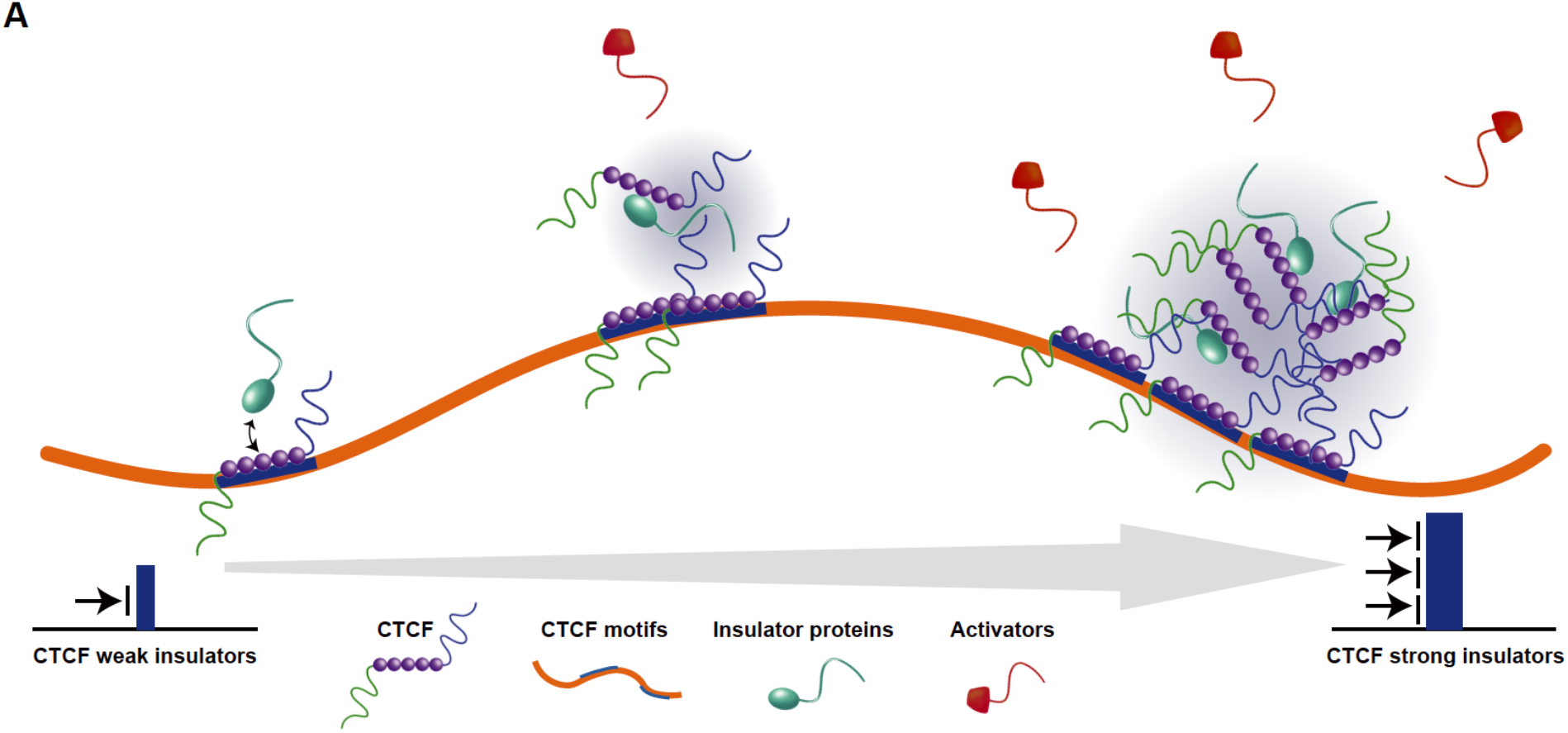
Spatial segregation model of CTCF insulation. CTCF binds its target motif, selectively interacts with insulator proteins, and preferentially avoids high densities of transcriptional activators. These properties lead to relatively high concentrations of insulator proteins and low concentrations of transcriptional activators at CTCF-bound positions, which will spatially segregate the communication of transcriptional activators between enhancers and promoters. As the binding strength of CTCF increases, its insulation capacity also increases. Note: the spatial segregation model was proposed based on the results of CTCF DBD, which needs further validation with the full-length endogenous CTCF in cells in the future.

